# A Geometric View of Signal Sensitivity Metrics in multi-echo fMRI

**DOI:** 10.1101/2022.02.23.481688

**Authors:** Thomas T. Liu, Bochao Li, Brice Fernandez, Suchandrima Banerjee

**Author notes:** Corresponding Author: Email address (Thomas T. Liu).

## Abstract

In multi-echo fMRI (ME-fMRI), two metrics have been widely used to measure the performance of various acquisition and analysis approaches. These are temporal SNR (tSNR) and differential contrast-to-noise ratio (dCNR). A key step in ME-fMRI is the weighted combination of the data from multiple echoes, and prior work has examined the dependence of tSNR and dCNR on the choice of weights. However, most studies have focused on only one of these two metrics, and the relationship between the two metrics has not been examined. In this work, we present a geometric view that offers greater insight into the relation between the two metrics and their weight dependence. We identify three major regimes: (1) a tSNR robust regime in which tSNR is robust to the weight selection with most weight variants achieving close to optimal performance, whereas dCNR shows a pronounced dependence on the weights with most variants achieving suboptimal performance; (2) a dCNR robust regime in which dCNR is robust to the weight selection with most weight variants achieving close to optimal performance, while tSNR exhibits a strong dependence on the weights with most variants achieving significantly lower than optimal performance; and (3) a within-type robust regime in which both tSNR and dCNR achieve nearly optimal performance when the form of the weights are variants of their respective optimal weights and exhibit a moderate decrease in performance for other weight variants. Insight into the behavior observed in the different regimes is gained by considering spherical representations of the weight dependence of the components used to form each metric. For multi-echo acquisitions, dCNR is shown to be more directly related than tSNR to measures of CNR and signal-to-noise ratio (SNR) for both task-based and resting-state fMRI scans, respectively.

## 1. Introduction

In multi-echo functional magnetic resonance imaging (ME-fMRI), time series data are acquired at various echo times (TE) and then typically combined to form a composite time series, formed as the weighted sum of the data from the different echoes. The potential benefits of ME-fMRI include the ability to increase sensitivity to functional changes and the capacity to more effectively distinguish BOLD-related functional signal changes from non-BOLD contributions (Kundu et al., 2017; Cohen et al., 2021).

In evaluating the performance of ME-fMRI methods, two measures of sensitivity have been typically used. These are (1) temporal signal-to-noise-ratio (tSNR) (Kundu et al., 2013; Puckett et al., 2018; Cohen et al., 2017; Heunis et al., 2021; Cohen et al., 2021) and (2) differential contrast-to-noise ratio (dCNR), also known as pseudo-temporal BOLD Sensitivity (ptBS) (Poser et al., 2006; Kettinger et al., 2016). In general, the contrast-to-noise ratio (CNR) of a functional time series is computed as the ratio of the functional signal change divided by the temporal standard deviation *σ*. For a single-echo acquisition, CNR can be expressed as the product of the normalized BOLD signal change multiplied by tSNR, defined as the temporal mean of the BOLD signal divided by *σ* (Parrish et al., 2000; Krüger and Glover, 2001). Thus, for an expected functional BOLD signal change (which depends on factors such as echo time, magnetic field strength, experimental design, and physiology), the CNR of a single-echo acquisition is maximized when tSNR is maximized.

For a multi-echo acquisition, the computation of CNR must take into account both the dependence of the BOLD signal change on echo time (TE) (Posse et al., 1999; Poser et al., 2006; Gowland and Bowtell, 2007) and the weight assigned to each echo. This results in an expression where the CNR can be expressed as the product of the expected change 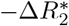 in effective transverse relaxation rate and dCNR, such that for a given change in the relaxation rate CNR is maximized when dCNR is maximized (see also Theory section below). To compute tSNR, the temporal mean of the composite time series is divided by its standard deviation. As discussed in the Theory section, dCNR and tSNR are not necessarily proportional and the relation between CNR and tSNR is not straightforward.

Prior work has considered the weight dependence of both metrics. In a seminal paper, Posse et al. (1999) considered various multi-echo weighting schemes and found that theoretical CNR was maximized with matched-filter weighting under the assumption that the noise is uncorrelated with equal variances across echoes. This weighting scheme has been adopted by a number of subsequent studies and referred to by a variety of names, such as optimal combination (Kundu et al., 2013), BOLD-signal weighted (Poser et al., 2006; Kettinger et al., 2016), and *T*_2_*-weighted (Heunis et al., 2021). Other weighting schemes have also been explored, such as a flat weighting in which all echoes have the same weight Posse et al. (1999); Poser et al. (2006); Gowland and Bowtell (2007). Poser et al. (2006) found that matched filter weighting provided slightly better dCNR than flat weighting, with both schemes performing slightly worse than a variation referred to as temporal BOLD sensitivity weighting. Using a different protocol, Kettinger et al. (2016) found that average dCNR values obtained with matched filter weighting were slightly higher than those obtained with either flat or temporal BOLD sensitivity weighting approaches, but concluded that all the weighting schemes provided similar group-level statistical performance. Recently, Heunis et al. (2021) found that various weighting schemes, including the matched filter weighting, provided comparable tSNR measures across the whole brain and selected regions of interest.

In this paper, we present a geometric perspective that provides further insight into the dependence of tSNR and dCNR on the weighting scheme. After reviewing the definitions of tSNR and dCNR, we examine the optimal solution for each metric and consider variants of the optimal solution, relating these variants to previously used weighting schemes. We then describe three major regimes of interest and show how insight into weight dependence in each regime can be gained by considering the geometric relation between the numerator and denominator terms of the metrics. Representative behavior of the metrics in each regime is demonstrated using example data from a multi-echo acquisition with three echoes, facilitating a 3D graphical view of the weight dependence. Preliminary versions of this work were presented in Liu et al. (2020,).

## 2. Theory

### 2.1 Review of definitions

In this section we briefly review the definitions of CNR, tSNR, and dCNR (Poser et al., 2006; Triantafyllou et al., 2005). To simplify the presentation, the spatial dependence of all quantities is assumed but not explicitly stated in the notation. For a given time series *S*, the contrast-to-noise-ratio is defined as 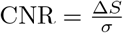 where Δ*S* is the signal change and *σ* is the standard deviation. For a single-echo acquisition this can be rewritten as

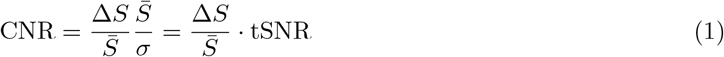

where 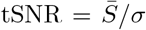 and 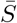 denotes the temporal mean. Thus, as noted in the Introduction, CNR for a single-echo acquisition is the product of the fractional signal change (equivalent to the percent signal change divided by 100) and tSNR.

For a multi-echo acquisition, the signal of interest is the weighted sum 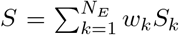, where *w*_*k*_ and *S*_*k*_ denote the weight and signal of the *k*th echo, respectively, and *N*_*E*_ is the number of echoes. From the basic definition of tSNR, we can write the multi-echo version as

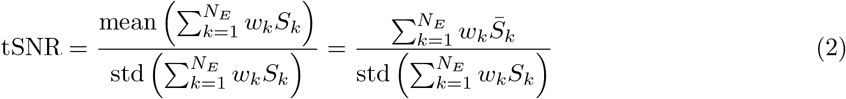

where 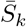 is the temporal mean of the *k*th echo.

To derive an expression for multi-echo CNR, we express the signal change term as 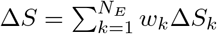 where Δ*S*_*k*_ is the signal change of the *k*th echo. As the fractional BOLD signal change for the *k*th echo can be well approximated as the product 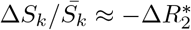. TE_*k*_ of the change in effective transverse relaxation rate 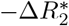 and echo time TE_*k*_, the signal change term may be written as 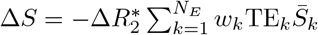 (Poser et al., 2006; Kundu et al., 2013). Dividing this term by the standard deviation of the signal yields

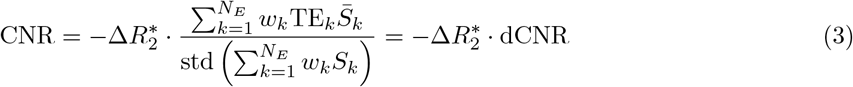

where differential CNR is defined as

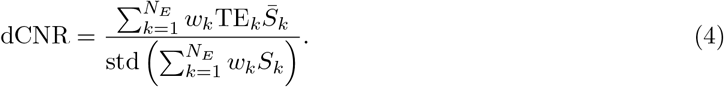

In comparing Equations 2 and 4, we can see that the expressions for tSNR and dCNR differ only in the addition of the echo time TE_*k*_ terms in the numerator for dCNR. For an arbitrary choice of echo times, tSNR is not necessarily proportional to either dCNR or CNR for a multi-echo acquisition. In the theoretical limiting cases where there is either just one echo (*N*_*E*_ = 1) or where there are multiple echoes with equal echo times (TE_*k*_ = TE), then dCNR = TE · tSNR. While the second limiting case cannot be achieved in practice, recently developed approaches can acquire echoes with a separation on the order of 1 ms and thereby closely approach the limiting case (Wang et al., 2019).

To facilitate geometric understanding, we rewrite tSNR and dCNR using matrix notation. We define **w** as the *N*_*E*_ × 1 vector of weights and **S** as a *N*_*E*_ × *N*_*T*_ matrix where the *i*th row contains the time series data from the *i*th echo, and *N*_*T*_ denotes the number of time points. With this notation **w**^*T*^ **S** represents the weighted sum of the signals and

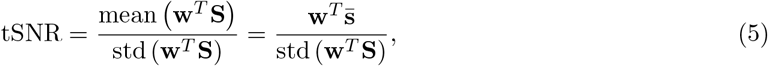

where 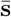 is the *N*_*E*_ × 1 vector consisting of the temporal means 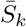. In a similar fashion, we can write

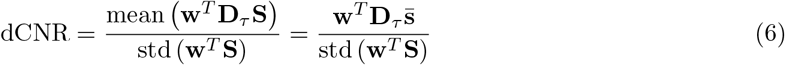

where 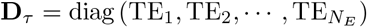 is the diagonal matrix comprised of the echo times. To support understanding of these expressions, expanded versions of Equations 5 and 6 for the case of *N*_*E*_ = 3 are provided in the Appendix.

### 2.2. Optimal Solutions

To maximize tSNR, we can maximize its square

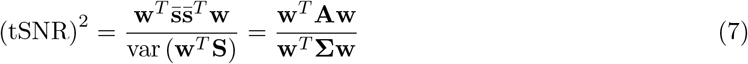

where 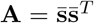 and **Σ** denotes the covariance matrix of **S**. The expression for (tSNR)^2^ has the form of a generalized Rayleigh quotient (GRQ) and attains a maximum value of 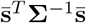 when the weight vector has the form of an optimal matched filter 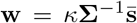, where *κ* is an arbitrary scalar (Brennan and Reed, 1973; Monzingo et al., 2011; Jarrett et al., 2017). From the form of the GRQ, it is straightforward to show that 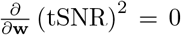 when **A** is a scalar multiple of **Σ**. Thus, we expect that tSNR will be relatively insensitive to the form of **w** when **A** ∼ **Σ**.

For dCNR, the corresponding GRQ is

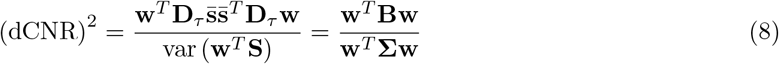

where 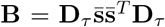. The GRQ attains its maximal value 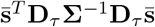 when 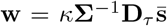. Furthermore, the GRQ and hence dCNR are both fairly insensitive to the choice of weights when **B** ∼ **Σ**. Since both tSNR and dCNR are invariant with respect to the value of *κ*, we will drop this scalar term for the remainder of the paper. To facilitate understanding of the GRQ expressions, expanded versions of Equations 7 and 8 for the case of *N*_*E*_ = 3 are provided in the Appendix.

Under certain conditions, previously proposed weighting schemes can achieve optimal tSNR or dCNR. For example, with the assumption that the random noise components have equal variance and are uncorrelated across the echoes (i.e. Σ = *σ*^2^**I**), the optimal weight vector for dCNR has the form 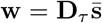 (i.e. 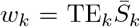), which is the BOLD Sensitivity weight solution examined in Kettinger et al. (2016). This is equivalent to the matched filter solution proposed in Posse et al. (1999) when the signal means exhibit 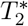 weighting (i.e. 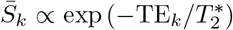), as would be the case for most ME-fMRI acquisitions. For tSNR maximization, the assumption Σ = *σ*^2^**I** leads to an optimal weight vector of the form 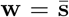 (i.e. 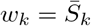), which is analogous to the exponential weighting considered by Posse et al. (1999).

If the noise is uncorrelated with unequal variances across echoes (i.e. 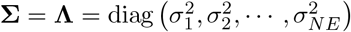), then the optimal dCNR weight vector has the form 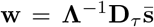 (i.e. 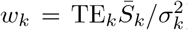). This solution is similar to but not equivalent to the temporal BOLD sensitivity weighting 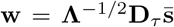 (i.e. 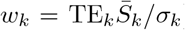) proposed in Poser et al. (2006) and further examined in Kettinger et al. (2016). For tSNR maximization, the optimal weight vector when **Σ** = **Λ** is 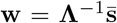 (i.e. 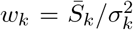), which is similar to but not equivalent to the tSNR weighting 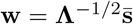 (i.e. 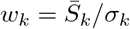) studied in Kettinger et al. (2016).

The various options described above for the weight vector are summarized in Table 1. The first four options consist of the tSNR optimal solution followed by three tSNR weight variants listed in order of decreasing similarity to the optimal solution. The last five options consist of the dCNR optimal solution followed by four dCNR weight variants listed in order of decreasing similarity to the optimal solution. In addition, we also consider a flat weighting of the echoes that has been examined in prior studies (Posse et al., 1999; Kettinger et al., 2016; Gowland and Bowtell, 2007).

**Table 1:** Description of weight vectors **w** where **s** denotes the vector of signal means, **Σ** denotes the noise covariance matrix, **Λ** is the diagonal matrix with noise variances along the diagonal, **D**_*τ*_ is the diagonal matrix composed of echo times on the diagonal, **I** is the identity matrix, and **1** is a column vector of 1’s. The 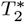 weighted solution (also referred to in the literature as matched filter or optimal combination) was originally proposed in Posse et al. (1999) and is equivalent to the BOLD Sensitivity weight examined in Kettinger et al. (2016) when the mean signal exhibits monoexponential 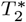 decay. Temporal BOLD sensitivity weighting was proposed by Poser et al. (2006) and tSNR weighting was proposed by Kettinger et al. (2016). The tSNR diagonal and dCNR diagonal weight vector are variants of the tSNR and dCNR optimal weight vectors, respectively, obtained by ignoring the off-diagonal elements of **Σ**. These are similar in form but not equivalent to the temporal BOLD sensitivity and tSNR weight vectors. The signal weighted solution is analogous to the exponential weighting considered by Posse et al. (1999).

| Method | Abbrev. | $\mathbf{w}$ | Comments |
| --- | --- | --- | --- |
| tSNR Optimal | topt | $\mathbf{\Sigma}^{-1}\bar{\mathbf{s}}$ | |
| tSNR Diagonal | tdg | $\mathbf{\Lambda}^{-1}\bar{\mathbf{s}}$ | tSNR optimal if $\mathbf{\Sigma}$ is diagonal |
| tSNR Weighted | tsnr | $\mathbf{\Lambda}^{-1/2}\bar{\mathbf{s}}$ | |
| Signal Weighted | swt | $\bar{\mathbf{s}}$ | tSNR optimal if $\mathbf{\Sigma} = \sigma^2\mathbf{I}$ |
| Flat | flat | $\mathbf{1}$ | |
| dCNR Optimal | dopt | $\mathbf{\Sigma}^{-1}\mathbf{D}_\tau\bar{\mathbf{s}}$ | |
| dCNR Diagonal | ddg | $\mathbf{\Lambda}^{-1}\mathbf{D}_\tau\bar{\mathbf{s}}$ | dCNR optimal if $\mathbf{\Sigma}$ is diagonal |
| Temporal BOLD Sensitivity | tBS | $\mathbf{\Lambda}^{-1/2}\mathbf{D}_\tau\bar{\mathbf{s}}$ | |
| BOLD Sensitivity | BS | $\mathbf{D}_\tau\bar{\mathbf{s}}$ | dCNR optimal if $\mathbf{\Sigma} = \sigma^2\mathbf{I}$ |
| $T_2^*$ Weighted | t2wt | $\mathbf{D}_\tau \exp(-\text{diag}(\mathbf{D}_\tau)/T_2^*)$ | dCNR optimal if $\mathbf{\Sigma} = \sigma^2\mathbf{I}$ and<br>$\bar{\mathbf{s}} \propto \exp(-\text{diag}(\mathbf{D}_\tau)/T_2^*)$ |

### 2.3. Relation to SNR in resting-state fMRI

As noted above, tSNR and dCNR can be used to predict the CNR that can be achieved when there is a task-related change in 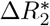 for single-echo and multi-echo task-based fMRI studies, respectively. For these studies, endogenous fluctuations in the BOLD signal are typically considered to be a noise component (in addition to other non-BOLD components) that can be distinguished from the task-based BOLD signal change. However for resting-state fMRI (rsfMRI) studies, the intrinsic BOLD fluctuations constitute the signal of interest while non-BOLD signal sources, such as physiological and thermal fluctuations, comprise the noise components (Liu, 2016).

The signal-to-noise ratio (SNR) in rsFMRI can be defined as rSNR = *σ*_*B*_*/σ*_*N*_ where *σ*_*B*_ and *σ*_*N*_ denote the standard deviations of the BOLD and non-BOLD components, respectively (Liu, 2013). For a single-echo acquisition, the BOLD term can be approximated as 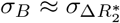 · TE · 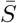 where 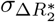 denotes the standard deviation of the fluctuations in 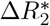. This allows us to write

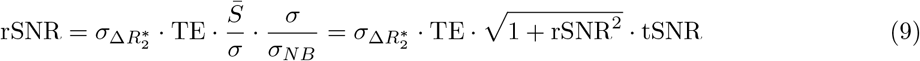

where the overall noise variance 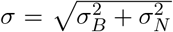 under the assumption that the BOLD and non-BOLD components are not correlated with each other. Rewriting this relation as

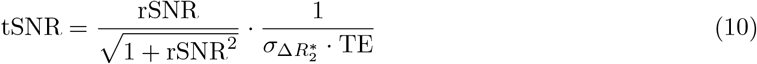

makes it easier to see that tSNR ∼ rSNR in the low rSNR regime (rSNR ≪ 1), whereas tSNR approaches the asymptote 1*/*(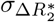 · TE) in the high rSNR regime (rSNR ≫ 1). The high rSNR asymptote can also be written in the form 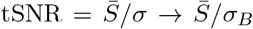, indicating that for a given value of *σ*_*B*_ the tSNR approaches 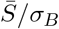 as the relative contribution of *σ*_*N*_ to *σ* is reduced (e.g. via noise reduction methods such as multi-echo independent components analysis (Kundu et al., 2013)). In the transition zone where rSNR ∼ 1, there is a monotonically increasing nonlinear relation between tSNR and rSNR that bridges across the linear and asymptotic zones.

For multiecho rsfMRI, we use matrix notation to write the relevant variance terms as 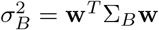 and 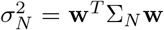 where Σ_*B*_ and Σ_*N*_ represent the covariance matrices for the BOLD and non-BOLD components, respectively. We can then write

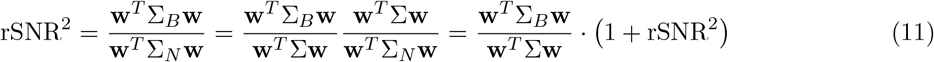

where the overall covariance matrix Σ = Σ_*B*_ +Σ_*NB*_, once again assuming that the BOLD and non-BOLD components are not correlated with each other. Using the previously stated approximation for BOLD component fluctuations 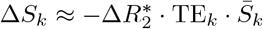, we obtain 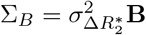 and write

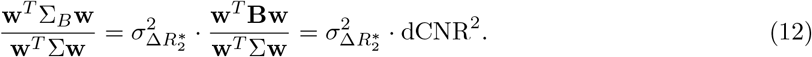

Inserting this into equation 11, rearranging terms, and taking the square root yields

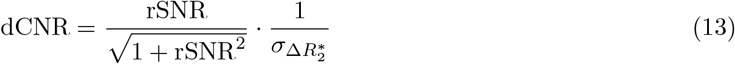

Note that the dependence of multi-echo dCNR on rSNR has the same form as that observed for single-echo tSNR within a scaling factor of TE.

### 2.4. Geometric View

In this section we consider three major regimes of interest with regards to the weight dependence of the sensitivity metrics.

1. **tSNR Robust Regime**: tSNR is relatively robust to the choice of weights with most variants achieving close to optimal performance, while dCNR shows a substantial dependence on the weights with most variants achieving significantly lower than optimal performance.
2. **dCNR Robust Regime**: dCNR is relatively robust to the choice of weights with most variants achieving close to optimal performance, while tSNR exhibits a substantial dependence on the weights with most variants achieving significantly lower than optimal performance.
3. **Within-type Robust Regime**: In this regime, both tSNR and dCNR are nearly optimal for weight variants that are similar to their respective optimal solution and exhibit moderately high performance for weight variants that are similar to the optimal solution for the other metric.

The tSNR and dCNR robust regimes are characterized by a high degree of similarity between their respective numerator matrices and the covariance matrix **Σ**, with **A** ∼ **Σ** for the tSNR robust regime and **B** ∼ **Σ** for the dCNR robust regime. Since **A** and **B** are both symmetric rank-one matrices formed as the outer products of a vector (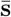 and 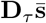, respectively) with its transpose, a high degree of similarity can be achieved when **Σ** is well-approximated by a rank-one matrix. This occurs when there is a dominant eigenvalue *λ*_1_ *λ*_*i* ≠1_ such that the covariance matrix may be approximated as 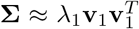, where **v**_1_ is the corresponding eigenvector.

To proceed, it is useful to examine the geometry of the numerator and denominator terms of the metrics in Equations 5 and 6. The numerator term is the inner product of the weight vector **w** with either the mean signal vector 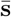 for tSNR or the TE-weighted mean signal vector 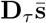 for dCNR. If the numerator term is normalized to have a maximum of 1.0, then the inner product as a function of weight can be written as either |cos (*θ*_*tSNR*_ (**w**))| or |cos (*θ*_*dCNR*_ (**w**))|, where *θ*_*tSNR*_ (**w**) and *θ*_*dCNR*_ (**w**) are the polar angles of **w** referenced to the vectors 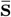 and 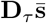 for tSNR and dCNR, respectively. Examples showing this cosine dependence are shown in Figure 1 panels (a) and (d) for a representative voxel, where the color of each point on the sphere indicates the value of the normalized numerator term given a unit-norm weight vector pointing in that direction. When the covariance matrix has a dominant eigenvalue, a similar argument can be used to show that the denominator term can be well approximated as 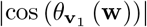 where 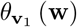 is the polar angle of **w** referenced to **v**_1_. Figure 1b shows an example of the corresponding surface plot for the representative voxel in which the dominant eigenvalue is 18.8 times larger than the second highest eigenvalue with a covariance fractional anisotropy (CFA) value of 0.96 (see Methods).

**Figure 1:**
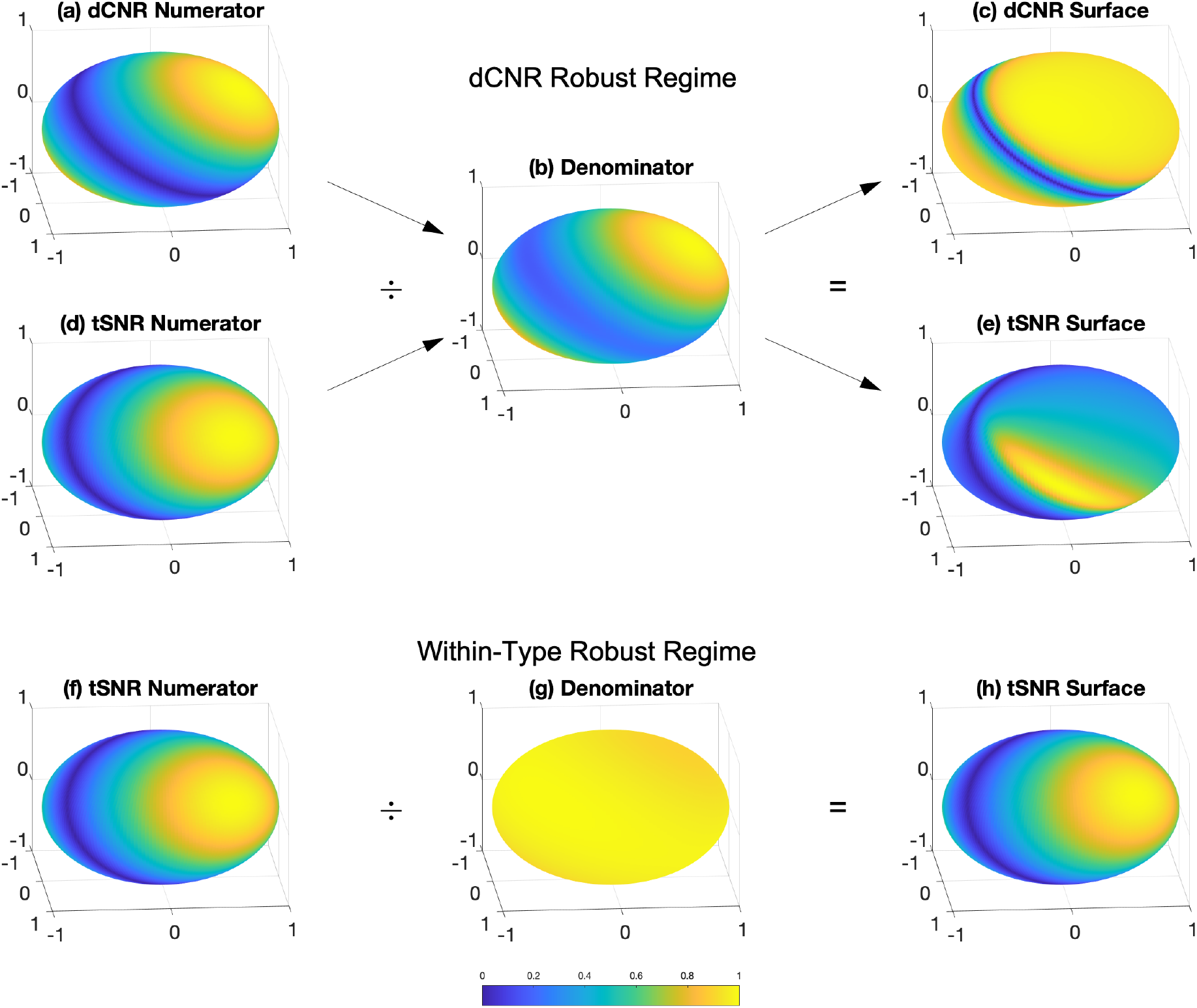
Geometric view of the dCNR robust regime and the within-type robust regime. The spherical surface plots indicate the value of the displayed quantities normalized by their maximum value, with each point on a sphere corresponding to a weight vector with unit norm. In the dCNR robust regime, the (a) dCNR numerator, the (d) tSNR numerator, and the (b) common denominator surface plots all exhibit a cosine dependence on the weights. The dCNR and tSNR surface plots are obtained by dividing the respective numerator terms by the common denominator terms. The robustness of the dCNR surface plot (panel c) reflects the relatively good alignment between the dCNR numerator and denominator surfaces. In contrast, the weight senstivity of the tSNR surface (panel e) reflects the misalignement between the tSNR numerator and denominator surfaces. In the within-type robust regime, the denominator surface (panel g) is fairly uniform, so that the tSNR surface (panel h) inherits the cosine dependence of the tSNR numerator surface (panel f).

The weight dependence of each metric reflects the ratio of the numerator and denominator surface plots. When the angle Δ*θ*_1_ between the dominant eigenvector **v**_1_ and the numerator reference vector (either 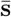 or 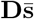) is relatively small, then the numerator and denominator surface plots will exhibit similar cosine dependencies on the weights and the corresponding metric is relatively insensitive to **w**. As an example, for the surface plots in Figure 1 panels (b) and (a) there is a relatively small angle Δ*θ*_1_ = 3.5^°^ between the reference vectors **v**_1_ and 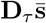, yielding a dCNR surface plot (panel c) that is relatively insensitive to the weights. However, since the angle between **v**_1_ and 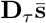 is small, this means that the angle between **v**_1_ and 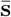 will be relatively large when the echo times span a range of values, as is typical for most multi-echo acquisitions. For the surface plots in panels (b) and (d), the angle between **v**_1_ and 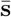 is Δ*θ*_1_ = 32.6^°^. The numerator and denominator terms exhibit cosine dependencies with significantly different directions, yielding a tSNR surface plot (panel e) that is highly sensitive to the weights. Thus, this example illustrates a key trade-off observed in the dCNR robust regime. When one metric (e.g. dCNR) is robust to the weights, the other metric (tSNR) will exhibit a high degree of dependence on the weights. The opposite tradeoff is observed in the tSNR robust regime, as described further in the Results section.

In the within-type robustness regime, the covariance matrix does not have a dominant eigenvalue and the denominator term shows a weak dependence on the weights. Figure 1g shows an example in which there is a fairly uniform spread of eigenvalues (normalized range of 1.00 to 1.25) leading to a relatively uniform denominator surface plot without a strong preferred direction. For this case, the tSNR metric shown in panel (h) primarily reflects the cosine dependence of the tSNR numerator term shown in panel (f). Furthermore, in this regime the covariance matrix can be roughly approximated **Σ** ∼ **I** by the identity matrix. With this approximation, the tSNR weight variants (tdiag, tsnr, swt) approach the tSNR optimal solution 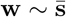 while the dCNR weight variants (ddg, tBS, BS, t2wt) approach the dCNR optimal solution 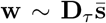. Thus, within each type there is a high degree of robustness to the weight selection when using a weight variant of that type (i.e. nearly optimal dCNR values obtained when using dCNR weight variants). Finally, because the metrics exhibit a cosine dependence on the weights, the performance when using a weight variant of the other type (e.g. decrease in dCNR when using a tSNR weight variant) can be approximated as the cosine of the angle between 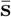 and 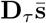. These concepts will be demonstrated in greater detail in the Results section.

### 2.5. Equivalence of normalized quantities

A specific case of the trade-off observed in the tSNR and dCNR robust regimes occurs when comparing tSNR computed using the dCNR optimal weight versus dCNR computed using the tSNR optimal weight. When these metrics are each normalized by their respective maximum values (obtained by using the tSNR and dCNR optimal weights, respectively), it can be shown that

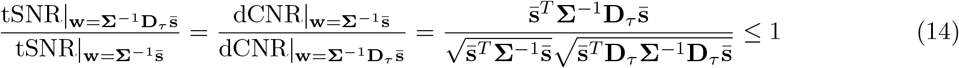

where the inequality follows from the Cauchy-Schwarz inequality (In other words, the normalized tSNR obtained with the dCNR optimal weight is equal to the normalized dCNR obtained with the tSNR optimal weight, and both quantities are less than or equal to 1.

## 3. Methods

To demonstrate the application of the geometric view, resting-state fMRI data (eyes-closed) were acquired under Institutional Review Board approval. One healthy male was scanned on a Discovery MR750 3T system (GE Healthcare) with a 32-channel receive head coil (Nova Medical) and a multi-echo multi-band EPI sequence with the following parameters: TR = 1.3 s; *θ* = 52^°^; FOV = 192 mm; matrix size = 64 × 64; 48 3 mm thick slices, multi-band factor = 3; TE=[12.2 30.1 48.0] ms, 277 reps; scan time = 6 min. Data were preprocessed using the meica.py script (v2.5b11) with 2nd order polynomial detrending, default despiking, and no smoothing (Kundu et al., 2013). A high-resolution anatomical scan (MPRAGE, flip angle = 8^°^, resolution = 1 mm^3^, FOV = 25.6 cm, TE = 2.92 ms, TR = 2500 ms, TI = 1060 ms, matrix size = 256×256×208) was acquired. Registration of the anatomical to the functional scan and estimation of tissue partial volume fractions were performed using AFNI and FSL (Cox, 1996; Smith et al., 2004).

For each voxel we computed values of tSNR and dCNR, using Equations 2 and 4 and the weight vectors (with sample means and covariances) listed in Table 1. We also computed the cosine similarities on a per-voxel basis between the the covariance matrix **Σ** and the numerator matrices **A** and **B** for tSNR and dCNR, respectively, using the six unique terms of each symmetric matrix. In addition, we computed the cosine similarity between **Σ** and the identity matrix. To characterize the spread of the eigenvalues of the covariance matrices, we adopted the fractional anisotropy (FA) measure used in diffusion tensor imaging (Basser and Pierpaoli, 1996) and computed this metric on a per-voxel basis using the covariance matrix eigenvalues. To make it clear that this metric does not refer to diffusion anisotropy, we will refer to this measure as covariance fractional anisotropy (CFA). Note that CFA values range from 0.0 to 1.0 with a value of 0.0 indicating that all the eigenvalues are equal and values approaching 1.0 indicating that the dominant eigenvalue is much greater than the other eigenvalues. Cosine similarity and CFA maps for five axial slices are shown in Figure 2.

**Figure 2:**
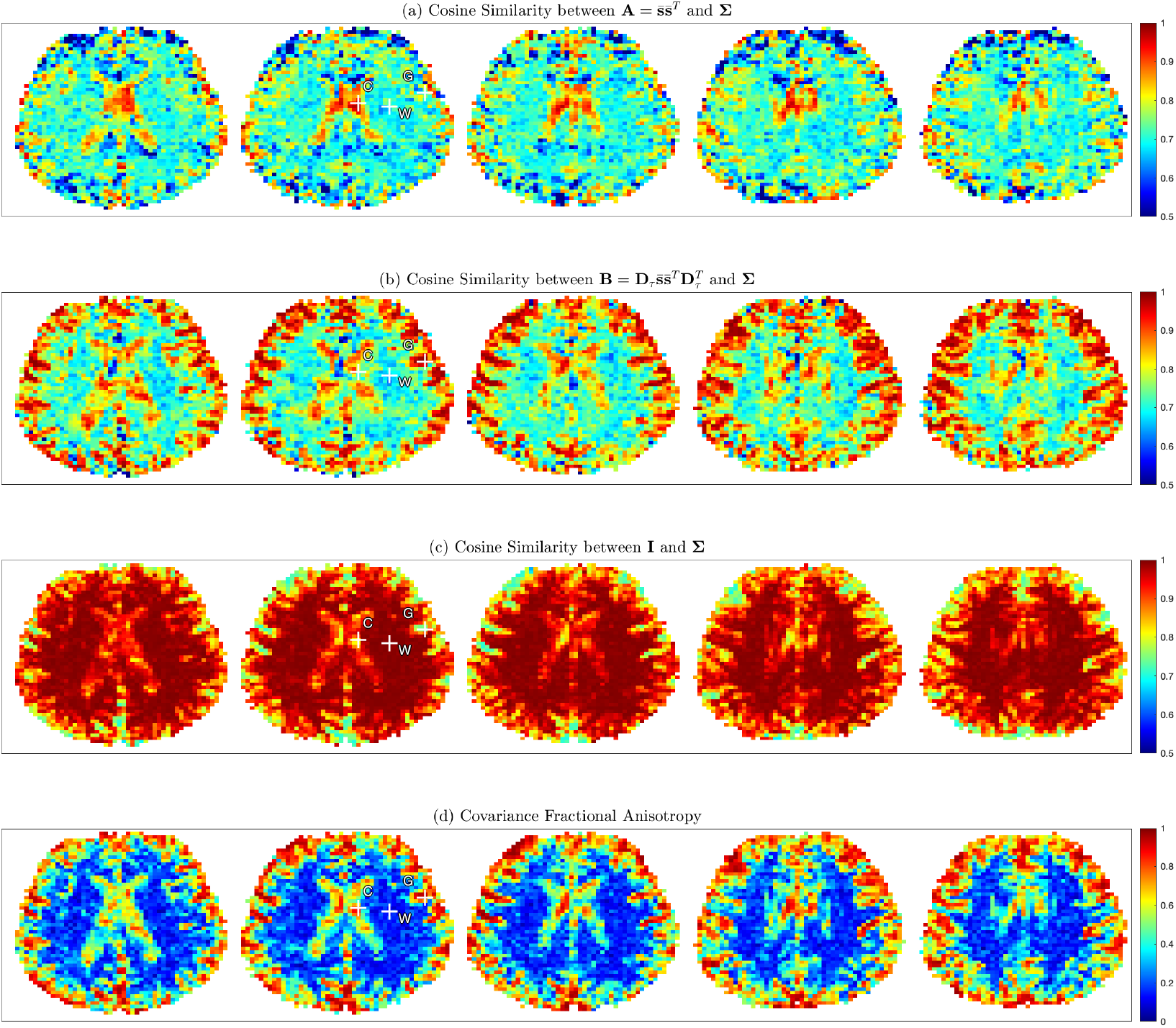
(a-c) Maps of cosine similarity between the covariance matrix **Σ** and (a) tSNR numerator matrix **A**, (b) dCNR numerator matrix **B**, and (c) the identity matrix **I**. Representative voxels for tSNR robust regime, dCNR robust regime, and within-type robust regime are marked with plus symbols and the letters C (CSF), G (gray matter), and W (white matter), respectively. (d) Covariance fractional anisotropy maps.

Based on the cosine similarity values, a representative voxel was chosen to illustrate each regime (indicated in Figure 2). A voxel (marked with cross and labeled with letter C) with high similarity (0.975) between **Σ** and the numerator matrix **A** was chosen to represent the tSNR robust regime, whereas a voxel (labeled with letter G) with high similarity (0.995) between **Σ** and **B** was chosen to represent the dCNR robust regime. Finally, a voxel (labeled with letter W) with high similarity (0.997) between **Σ** and the identity matrix **I** was chosen to represent the within-type robust regime. The cerebral spinal fluid (CSF), gray matter, and white matter partial volume fractions for voxels C, G, and W were 1.0, 0.54, and 1.0, respectively. The remaining volume fraction (0.46) in voxel G was assigned to CSF.

## 4. Results

In this section, we first examine in detail the characteristics of the representative voxel for each regime and show how the geometric view facilitates an understanding of the observed performance. We then consider the application of the geometric view to brain-wide data.

### 4.1. tSNR robust regime

As an example of behavior in the tSNR robust regime, Figure 3a shows tSNR and dCNR values as a function of weight variant for representative voxel C. For all panels in the figure, the tSNR and dCNR values are normalized by their corresponding maximum values of 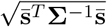 and 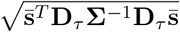, which are obtained using the respective optimal weight vectors. In this regime, the tSNR values are relatively robust to the choice of weight vector (ranging from 0.97 to 1.0), with the exception of the noticeably lower value of 0.70 for the dCNR optimal weight. In contrast, the dCNR values exhibit a greater degree of dependence on the choice of weight with values ranging from 0.53 to 1.0. The dotted black line highlights the equivalence stated previously in Equation 14, i.e. the normalized tSNR when using the optimal dCNR weight is equal to the normalized dCNR when using the optimal tSNR weight. (This equivalence is also highlighted for the other regimes shown in rows b and c).

**Figure 3:**
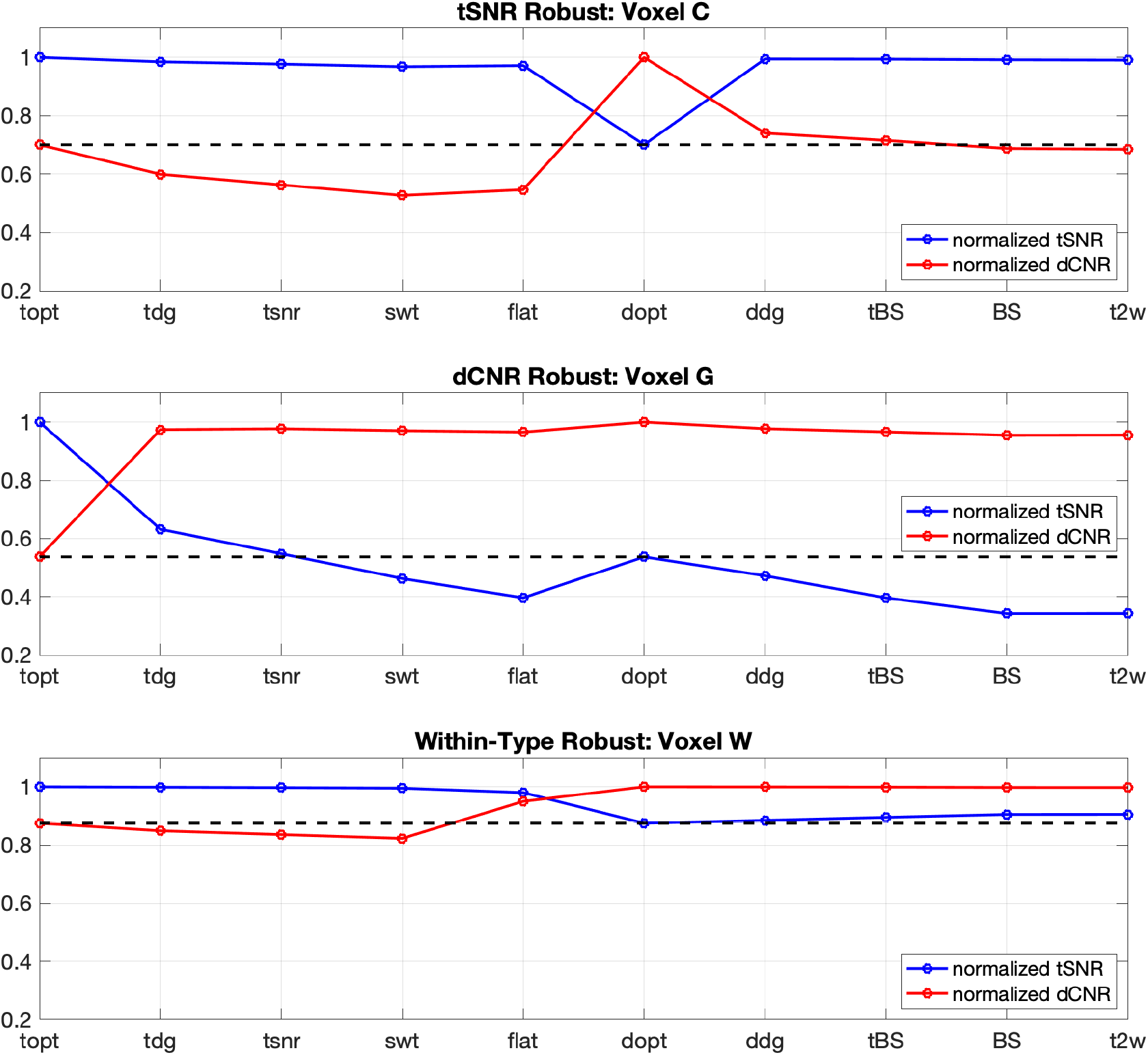
Normalized tSNR and dCNR values plotted versus weight vector variant for the representative voxels indicated in Figure 2. Weight vector variants are labeled with the abbreviations listed in Table 1. The black dotted lines are provided to point out that the normalized tSNR and dCNR values obtained when using dCNR and tSNR optimal weights, respectively, have the same value.

A geometric view of the tSNR robust regime is provided in the left column of Figure 4. In this figure the first and last rows show the absolute values of the normalized tSNR and dCNR values, respectively, plotted on the surface of a sphere, where each point of the sphere corresponds to a weight vector with unit norm. The tSNR and dCNR weight variants are indicated with brown squares and magenta circles, respectively, and the flat weight vectors are indicated by green diamonds. The second and fourth rows show the absolute values of the tSNR 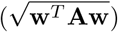 and dCNR 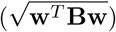 numerator terms, respectively, while the third row show the common denominator term 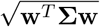. As described in detail in the figure caption, the symbols on the surface indicate the positions of the different weight variants.

**Figure 4:**
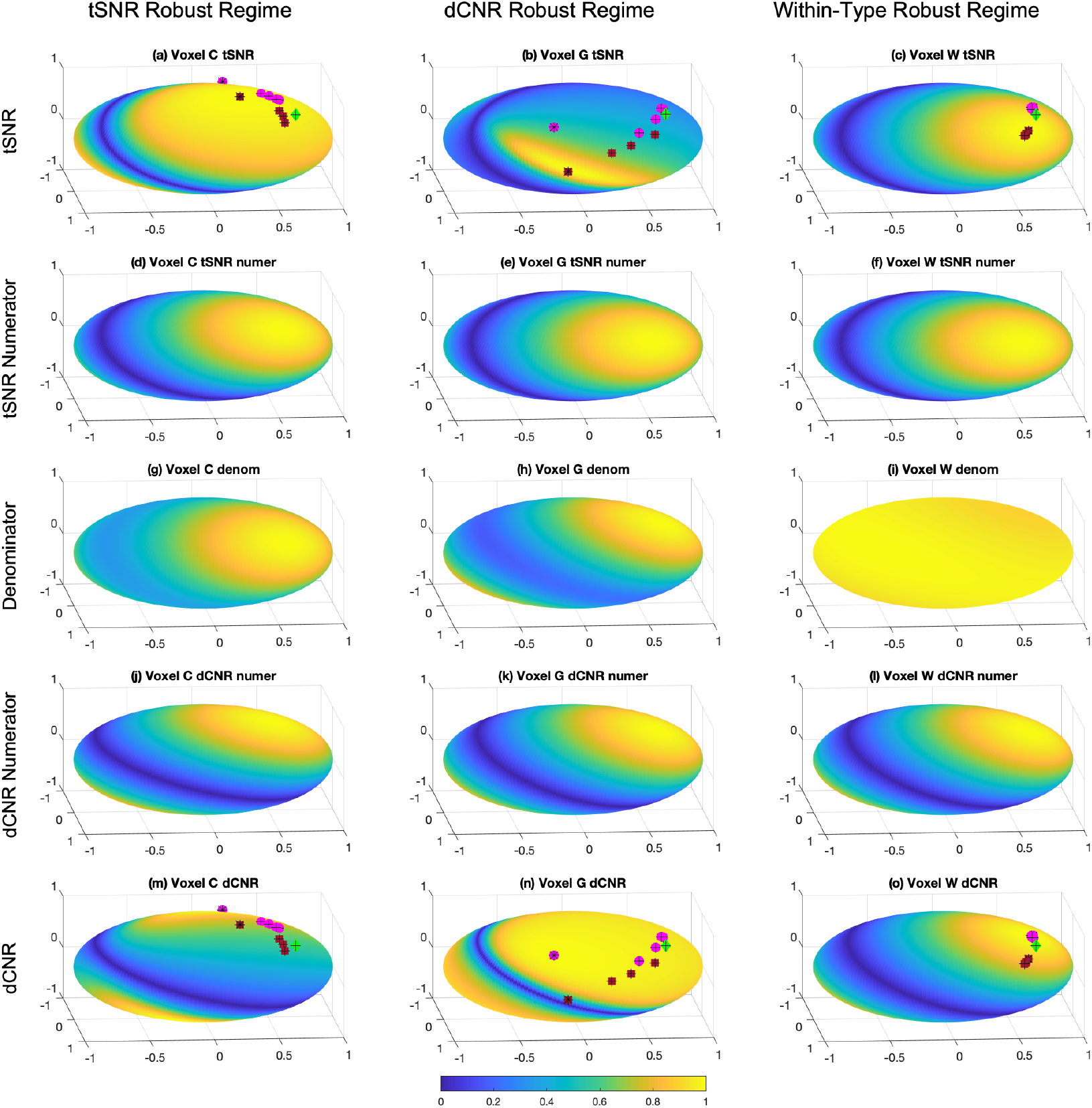
Spherical surface plots showing the weight dependence of (a-c) tSNR, (d-f) tSNR numerator, (g-i) denominator, (j-l) dCNR numerator, and (m-o) dCNR for representative voxels in the tSNR robust regime (voxel C; left column), dCNR robust regime (voxel G; center column), and the within-type robust regime (voxel W; right column). All quantities are shown as absolute values and normalized by their maximum value. The brown squares indicate tSNR weight variants (topt, tdg, tsnr, swt), the magenta circles indicates dCNR weight variants (dopt, ddg, tBS, BS, t2wt), and the green diamonds indicate the flat weight vectors. The optimal tSNR and dCNR weights are indicated with black asterisks while the other variants have black crosses.

Focusing on the left column for voxel C, both the tSNR and dCNR numerator surfaces (panels d and j) exhibit the cosine dependence described in Section 2.4. In addition, the denominator surface (panel g) also exhibits a cosine dependence because the covariance matrix has a dominant eigenvalue that is 7.1 times larger than the second highest eigenvalue, with a corresponding CFA value of 0.86. As the angle between the associated dominant eigenvector and the vector 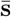 is relatively small (Δ*θ* = 6.3^°^), the tSNR numerator and denominator surfaces show a similar weight dependence, and the ratio of the two surfaces leads to a tSNR surface plot (panel a) that is relatively insensitive to the weights, consistent with the behavior shown by the green line in Figure 3a. In contrast, the principal directions of the dCNR numerator and denominator are not well aligned (Δ*θ* = 33.4^°^), so that the ratio of these two terms results in a dCNR surface plot (panel m) that is sensitive to **w**, consistent with the behavior shown by the red line in Figure 3a.

### 4.2. dCNR robust regime

To demonstrate behavior in the dCNR robust regime, Figure 3b shows tSNR and dCNR values as a function of weight variant for representative voxel G. The dCNR values are relatively robust to the choice of weight variant (ranging from 0.96 to 1.0), with the exception of the noticeably lower value of 0.54 for the tSNR optimal weight (which is equal to the tSNR value obtained with the dCNR optimal weight, as indicated by the black dotted line). In contrast, the tSNR values exhibit a high degree of weight sensitivity with values ranging from 0.34 to 1.0.

The geometric view of this regime was previously described in Section 2.4 and Figure 1. For completeness, it is presented in a slightly different format in the middle column of Figure 4. The dCNR numerator (panel k) and denominator (panel h) exhibit similar cosine dependencies, reflecting the relatively small angle (Δ*θ* = 3.5^°^) between 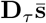 and the dominant eigenvector of the covariance matrix.

As a result, the ratio results in a dCNR surface plot (panel n) that is relatively insensitive to **w**. In contrast, the principal directions of the tSNR numerator (panel e) and denominator surface plots are not well aligned (Δ*θ* = 32.6^°^), so that the ratio of results in the tSNR surface plot (panel b) that is highly dependent on **w**.

### 4.3. Within-type robust regime

As an example of behavior in the within-type robust regime, Figure 3c shows tSNR and dCNR values across weight variants for voxel W. The normalized tSNR values for the tSNR optimal weight variants (topt, tdiag, tsnr, swt) are within 0.005 of the maximum value of 1.000, reflecting the fact that all of the variants approach the optimal solution 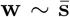 when **Σ** ∼ **I**. Similarly, the dCNR values for the dCNR optimal weight variants (dopt, ddg, tBS, BS, t2wt) are within 0.003 of the maximum value of 1.000, reflecting the fact that all of these variants approach the optimal solution 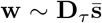. In considering the behavior of the other points, we note that both the tSNR values using the dCNR weight variants (ddg, tBS, BS, and t2wt) and the dCNR values using the tSNR weight variants (tdg, tnsr, and swt) lie fairly close to the dotted black equivalent value line (corresponding to tSNR with optimal dCNR weight (dopt) and dCNR with optimal tSNR weight (topt)).

The geometric view of tSNR in this regime was previously described in Section 2.4 and Figure 1. In the right column of Figure 4 we expand upon the prior description. Due to the fairly uniform spread of the covariance matrix eigenvalues (normalized range of 1.0 to 1.3, CFA = 0.12), the denominator surface (panel i) is fairly uniform. As a result, both the tSNR (panel c) and dCNR (panel o) surface plots reflect the cosine dependence of their respective numerator terms (panels f and l). In contrast to the spread in weight variant positions observed in the other regimes, the weight variants for voxel W are fairly tightly clustered within each variant type. As noted above, the tSNR and dCNR weight variants approach their respective optimal solutions (indicated with asterisks) in this regime, so that for tSNR in panel (c) the tSNR weight variants appear in the brown cluster, while for dCNR in panel (o) the dCNR weight variants appear in the magenta cluster. Each of these surfaces also contains a secondary cluster, corresponding to tSNR values when using dCNR weight variants (magenta cluster in panel c) and dCNR values when using tSNR weight variants (brown cluster in panel o). These are offset from the optimal clusters by an angle that is roughly the difference 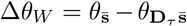 between the angles of 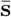 and 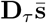. The cosine of this difference angle cos (Δ*θ*_*W*_) = 0.87 indicates reflects the reduced performance of the secondary cluster as compared to the maximum value of 1.0 obtained within the optimal cluster. In addition, it approximates the equivalent value shown by the black dotted line in Figure 3c, because the expression in Equation 14 is well-approximated by cos (Δ*θ*_*W*_) when **Σ** ∼ **I**.

### 4.4. Application to brain-wide data

To demonstrate the application of the geometric view to brain-wide data, we assigned each voxel to the regime that exhibited the highest corresponding cosine similarity value, where the similarities between the covariance matrix **Σ** and the matrices **A, B**, and **I** correspond to the tSNR robust regime, the dCNR robust regime, and the within-type robust regime, respectively. For example, a voxel for which the cosine similarity was highest between **Σ** and **B** was assigned to the dCNR robust regime. This process resulted in the assignment of 3.6% of the voxels to the tSNR robust regime, 19.2% to the dCNR robust regime, and 77.2% to the within-type robust regime. In the tSNR robust regime, the mean cerebral spinal fluid (CSF), gray matter (GM), and white matter (WM) partial volume fractions were 0.51, 0.35 and 0.14, respectively. For the dCNR robust regime, the mean partial volume fractions were 0.35 (CSF), 0.46 (GM), and 0.19 (WM), while for the within-type robust regime, the mean fractions were 0.09 (CSF), 0.33 (GM), and 0.58 (WM), respectively.

Figure 5 shows double-sided violin plots of the distribution within each regime of the normalized tSNR (blue distributions) and dCNR (red distributions) values for each weight variant, with the median values for each metric and weight indicated by the solid lines and open circles. To provide a complementary qualitative perspective, Figure 6 shows normalized tSNR and dCNR maps for the same five representative slices used in Figure 2. In Figure 5, the distributions for the tSNR optimal and dCNR optimal weight vectors are singular (e.g. all values equal to 1.0) for the tSNR and dCNR robust regimes, respectively. The equivalence stated in Equation 14 is demonstrated by the match between the shapes of the dCNR distributions (red) for the tSNR optimal weight (topt) and the tSNR distributions (blue) for the dCNR optimal weight (dopt). Furthermore, in Figure 6, the tSNR maps for weight option dopt are identical to the dCNR maps for weight option topt. The tradeoff between dCNR and tSNR is further demonstrated by the noticeably lower median values for these distributions in Figure 5 and evident in the wide spread of values in the corresponding maps (i.e. tSNR maps for dopt and dCNR maps for topt) shown in Figure 6. Due to the unique behavior of the distributions and maps obtained for the tSNR and dCNR optimal weights, the remainder of this section will focus on the performance of the other weight variants.

**Figure 5:**
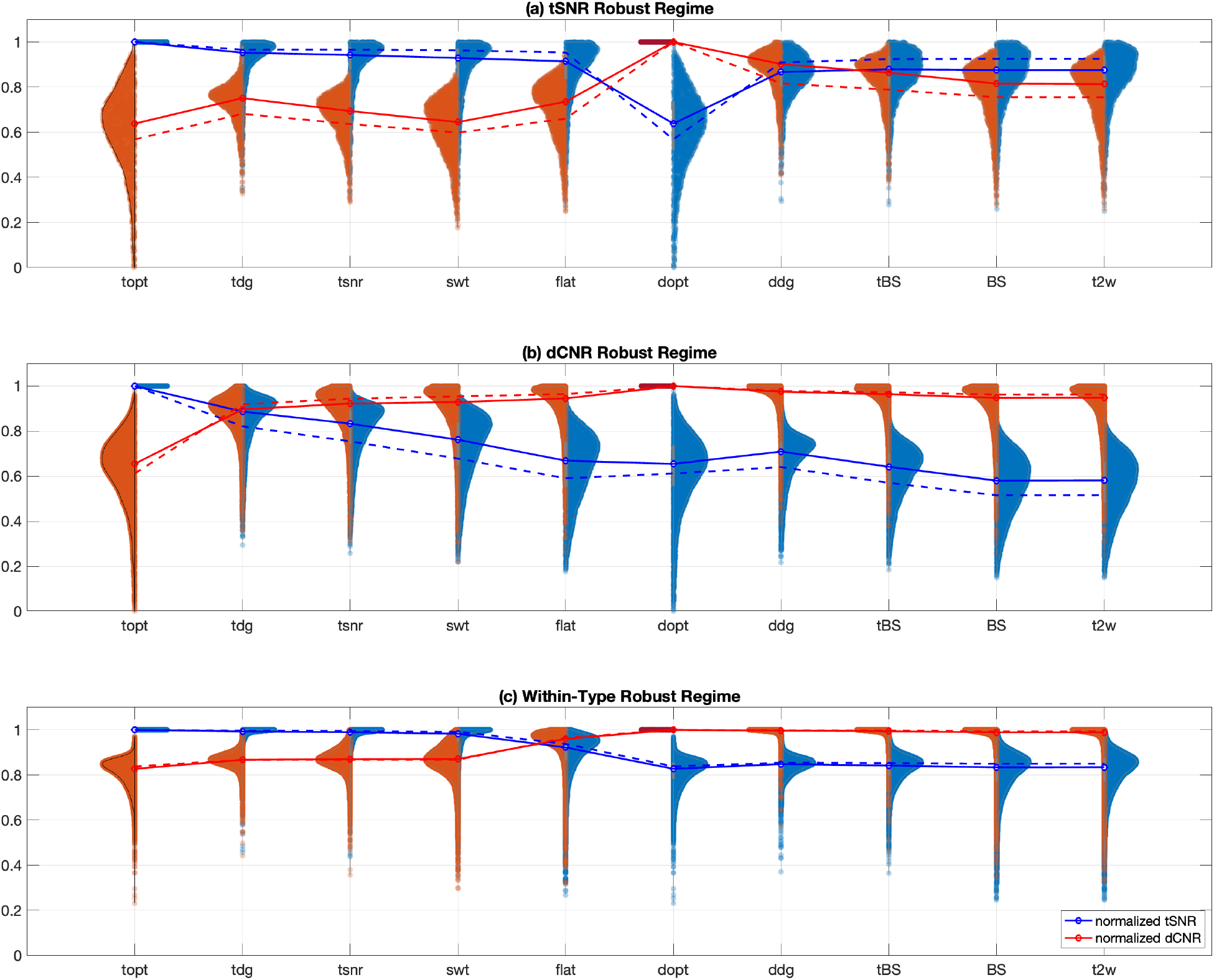
Two-sided violin plots showing the distributions of normalized tSNR (red) and dCNR (blue) values for each of the three regimes. Solid lines and circles indicate the median values of each distribution. Dashed lines indicate the median values obtained when a threshold of 0.95 on cosine similarity values is applied.

**Figure 6:**
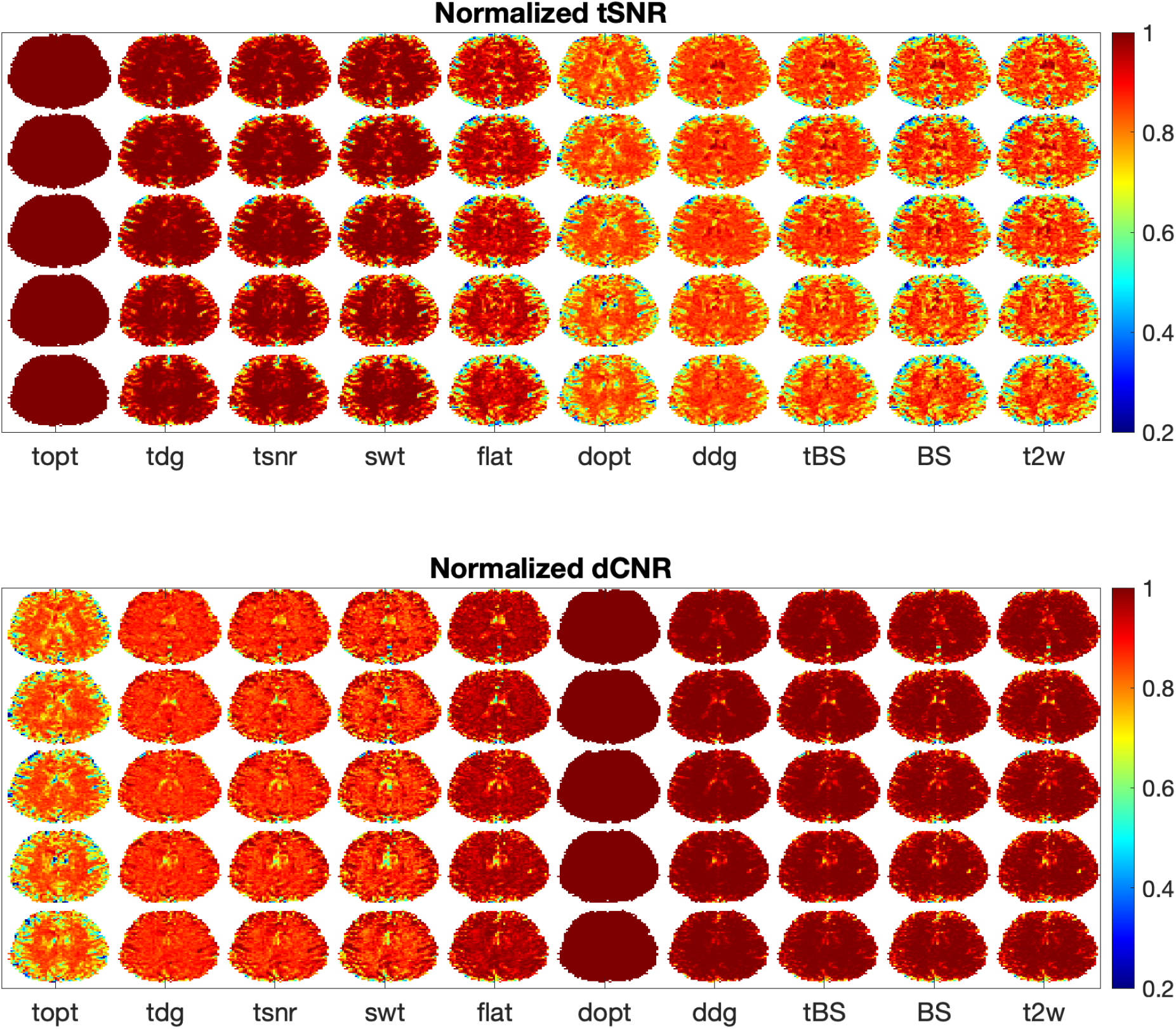
Maps of normalized tSNR and dCNR for 5 slices and 10 different weight vectors. Columns are labeled using the abbreviations listed in Table 1. Maps are normalized by the values obtained using their respective optimal weights, such that the maximum value is 1.0. Thus, the tSNR and dCNR maps obtained using the tSNR and dCNR optimal weights (topt and dopt), respectively, are both uniformly equal to 1.0, while maps using other weight vectors have values less than 1.0.

In the tSNR robust regime, the median tSNR values for the eight non-optimal weight variants range from 0.87 to 0.95 while the median dCNR values show greater variability, ranging from 0.58 to 0.89. In the normalized maps, this corresponds to small spots of high tSNR values in the ventricles (most clearly seen in the dCNR weight variants ddg, tBS BS, and t2w) and relatively lower dCNR values in these spots. For the dCNR robust regime, the median dCNR values range from 0.90 to 0.98, while the median tSNR values exhibit variations over a larger range from 0.65 to 0.90. From a qualitative perspective, the variation of dCNR values in the cortical regions is visibly lower than the variation observed for tSNR values. In the within-type robust regime, the median tSNR values for the tSNR weight variants range from 0.98 to 0.99 while the dCNR values for the dCNR weight variants range from 0.99 to 1.00. This is consistent with the relatively uniform and high tSNR and dCNR values observed in white matter regions for their respective weight variants.

Since the assignment of voxels to the regimes was based solely on the maximum cosine similarity observed for each voxel, each regime will contain voxels spanning a range of cosine similarity values. We also considered performance when restricting the analysis to include only voxels with higher cosine similarity values within each regime. The dashed-lines in Figure 5 show the median values obtained after applying a threshold of 0.95 to the cosine similarity values. There was a slight increase in the median values, and the effect was most pronounced for the tSNR values in the tSNR robust regime, with median tSNR values ranging from 0.91 to 0.97, as compared to the range of 0.87 to 0.95 obtained before thresholding. The effect of thresholding was much smaller for the other regimes. The partial volume fractions after thresholding for the tSNR robust regime were 0.67 (CSF), 0.24 (GM), and 0.09 (CSF); for the dCNR robust regime: 0.43 (CSF), 0.44 (GM), and 0.13 (CSF); and for the within-type robust regime: 0.04 (CSF), 0.29 (GM), and 0.67 (WM).

### 4.5. Factors Affecting Performance

To explain the modest increase in performance obtained with thresholding, we first note that the tSNR and dCNR distributions exhibit central modes with long tails towards smaller metric values, consistent with the occurrence of a range of lower values in the normalized maps. Voxels with lower cosine similarity values deviate more from the assumptions for robustness and are thus expected to exhibit lower metric values. This behavior is demonstrated in the two-dimensional scaled histograms of Figure 7, which display the probability and metric values in each regime as a function of cosine similarity and CFA for the signal weighted (swt) and BOLD sensitivity (BS) weight variants. The probabilities (indicated by the height of the bars) are highest around the median cosine similarity values of 0.92, 0.93, and 0.98 for the tSNR robust, dCNR robust, and within-type robust regimes, respectively. For all regimes, the normalized tSNR and dCNR values (indicated by the colors of the bars) tend to decrease with the relevant cosine similarity metric, with a strong monotonic relation observed for the within-type robust regime. Furthermore, the cosine similarity metric and CFA are tightly linked in the within-type robust regime, since both reflect the degree of anisotropy in the covariance matrix. Thresholding at 0.95 has very little effect on the metric values in the within-type robust regime because the median cosine similarity value of 0.98 is higher than the threshold.

**Figure 7:**
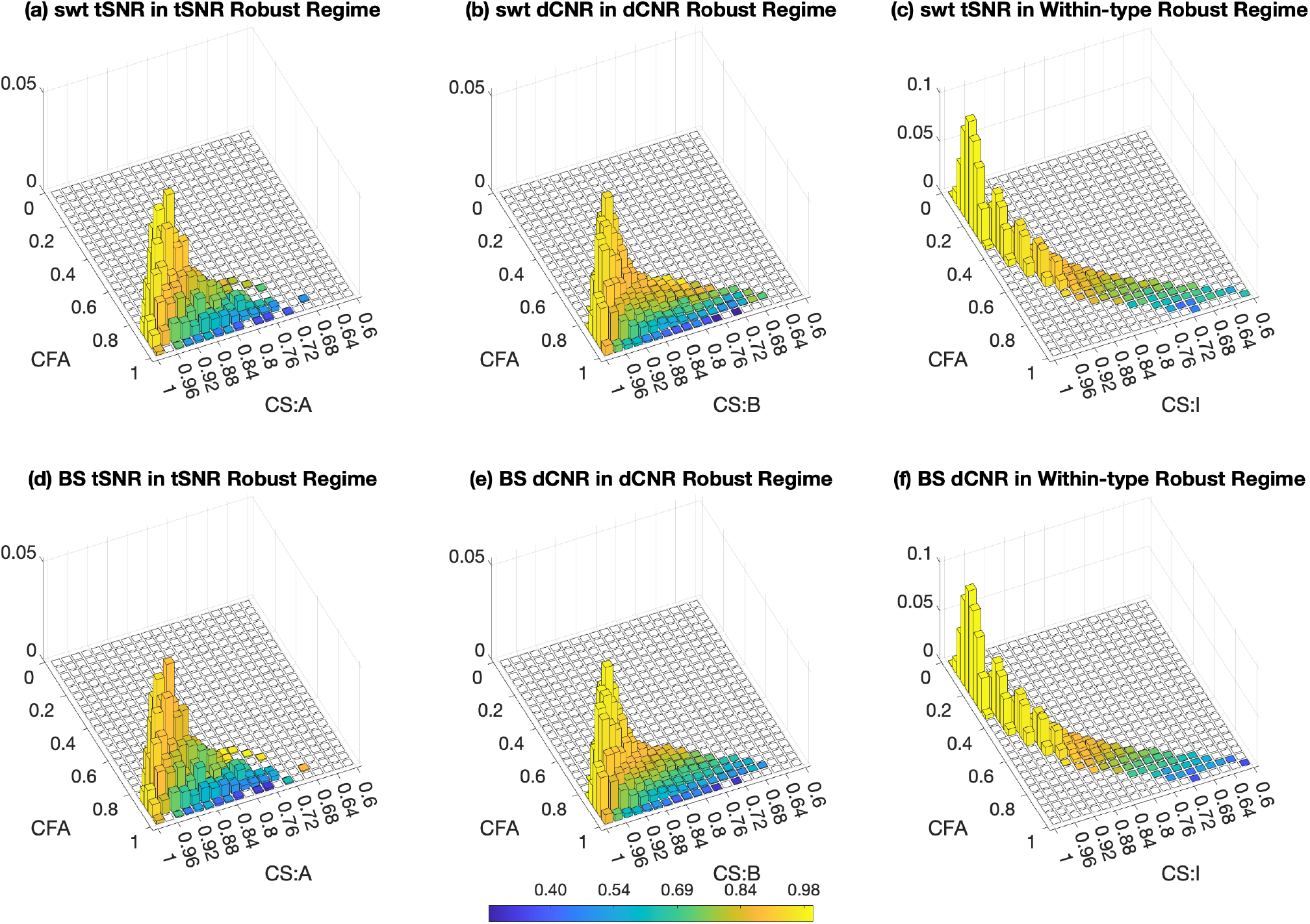
Scaled two-dimensional histograms showing color-coded normalized tSNR and dCNR values and probability of occurrence (height of bars) as a function of cosine similarity (CS) and covariance fractional anisotropy (CFA). Column 1 (panels a,d) shows normalized tSNR values of voxels in the tSNR robust regime for weight variants swt and BS as a function of CFA and cosine similarity (CS:A) between **Σ** and **A**. Column 2 (panels b,e) shows normalized dCNR values of voxels in the dCNR robust regime for weight variants swt and BS as a function of CFA and cosine similarity (CS:B) between **Σ** and **B**. Column 3 (panels c,f) shows normalized tSNR and dCNR values in the within-type robust regime for weight variants swt and BS, respectively, as a function of CFA and cosine similarity (CS:I) between **Σ** and **I**.

The metrics in the tSNR and dCNR robust regimes exhibit a dependence on CFA that is independent of cosine similarity values, with small tSNR and dCNR values observed for higher CFA values even when the cosine similarity value is high. In the tSNR and dCNR robust regimes, thresholding has some effect due to the lower median cosine similarity values in these regimes, but the effect is limited by the presence of small metric values occurring for high cosine similarity and CFA values. These values result in the persistence of long tails in the distributions even when thresholding based on cosine similarity is applied. In the next few paragraphs, we take a deeper look at the dependence of the metrics on cosine similarity and CFA in the tSNR and dCNR robust regimes.

As discussed in Section 2.4, the metrics in the tSNR and dCNR robust regimes are relatively insensitive to the choice of weight when the angle Δ*θ*_1_ between the dominant eigenvector **v**_1_ and the numerator reference vector (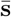 or 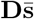) is relatively small. This is consistent with a high degree of similarity between the covariance matrix **Σ** and the numerator matrices **A** and **B** for the tSNR and dCNR robust regimes, respectively. As the similarity decreases, the angle Δ*θ*_1_ will tend to increase, resulting in greater weight sensitivity of the tSNR and dCNR surface plots. An example of this effect is shown in Figure 8, where the left and right columns show the denominator, dCNR numerator, and dCNR surfaces for two voxels with similar CFA values (0.83 and 0.84) but different cosine similarity (between **Σ** and **B**) values of 0.96 and 0.87, respectively, and associated angles Δ*θ*_1_ of 9.7 and 19.6 degrees. As expected the dCNR surface (panel i) associated with the larger angle exhibits a greater degree of weight dependence due to the larger mismatch in the directions of the denominator and numerator surfaces.

**Figure 8:**
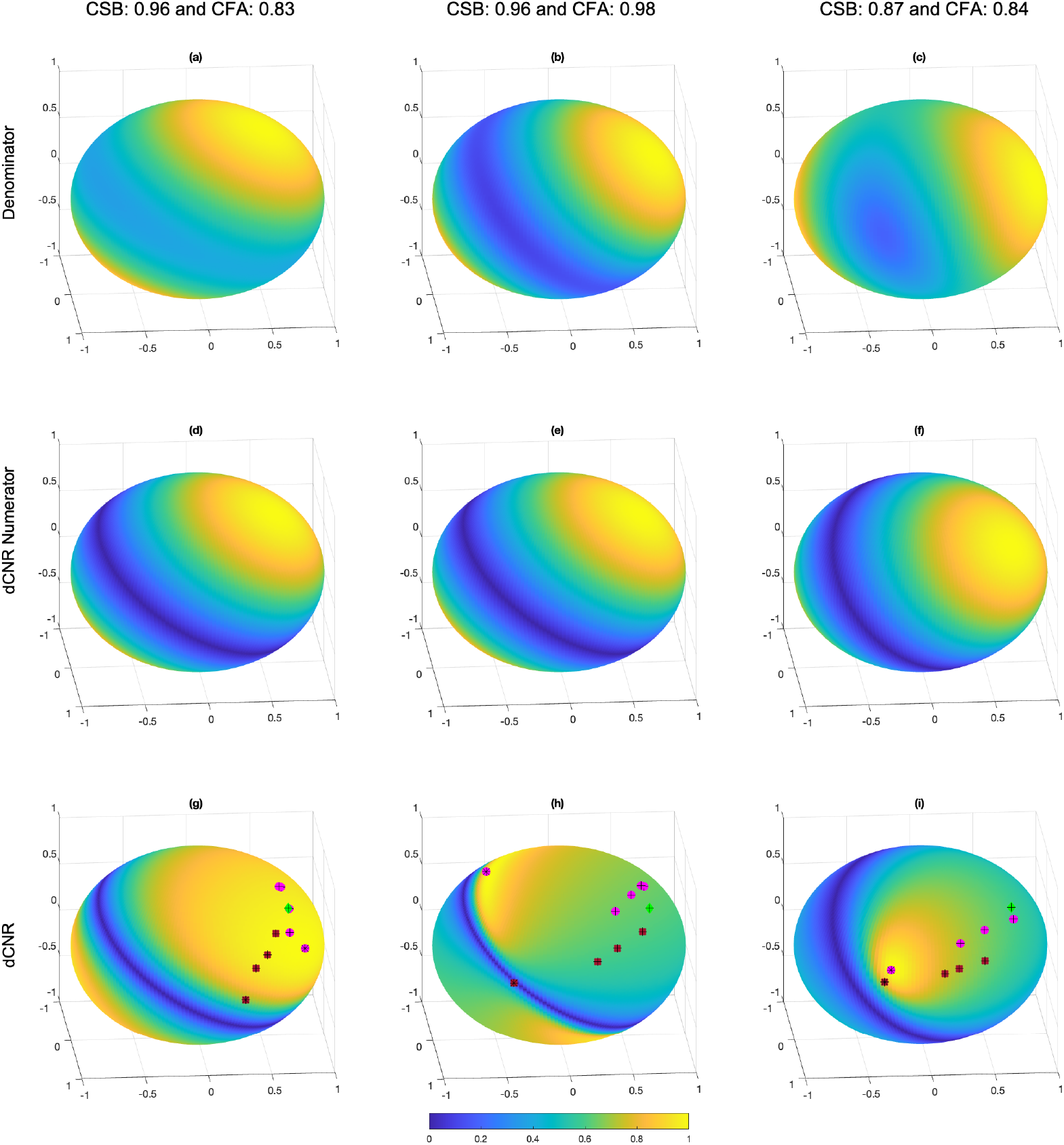
Spherical surface plots showing the weight dependence of (a-c) denominator, (d-f) dCNR numerator, and (g-h) dCNR for a voxel (left column) with high cosine similarity (CSB) between **Σ** and **B** and moderate covariance fractional anisotropy (CFA); a voxel (middle column) with high CSB and high CFA; and a voxel (right column) with moderate CSB and CFA values. All quantities are shown as absolute values and normalized by their maximum value. The brown squares indicate tSNR weight variants (topt, tdg, tsnr, swt), the magenta circles indicates dCNR weight variants (dopt, ddg, tBS, BS, t2wt), and the green diamonds indicate the flat weight vectors. The optimal tSNR and dCNR weights are indicated with black asterisks while the other variants have black crosses.

The observed dependence on CFA primarily reflects two effects: (1) an increase in weight dependence as CFA approaches 1.0 and (2) an increase in the angular distance between the optimal weight vector and other weight variants. To understand the first effect it is useful to recall that the sensitivity metrics are computed as the ratio of a numerator term that exhibits a pure cosine dependence on the weights and a denominator term that exhibits an approximate cosine dependence on the weights. In the limit where CFA = 1.0 there is only one non-zero eigenvalue and the denominator term exhibits a pure cosine dependence that will pass through zero for weights that are orthogonal to the dominant eigenvector. Division by this zero term would result in a singularity in the ratio. In practice, as CFA approaches 1.0 the minimum in the approximate cosine-like denominator term will be close to zero, resulting in a narrow peak in the ratio term and a sensitivity surface that is highly dependent on the weights. An example of this is shown in Figure 8, where the leftmost and middle columns show the denominator, dCNR numerator, and dCNR surfaces for two voxels with the same cosine similarity (between **Σ** and **B**) values (both 0.96) but different CFA values of 0.83 and 0.98. The denominator surface (panel b) for the voxel with higher CFA exhibits lower minimum values (darker blue colors) resulting in a more highly peaked and variable dCNR surface, as shown in panel h.

We can get a sense of the effect of CFA on the angular distance between the weights by considering the angle between the dCNR optimal weight vector 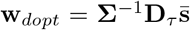 and the BOLD sensitivity weight variant 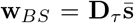. Using the eigenvector expansion 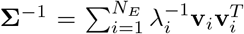, we can write 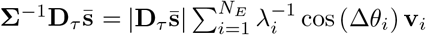 where Δ*θ*_*i*_ denotes the angle between 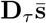 and the *i*th eigenvector **v**_*i*_. For high cosine similarity values in the dCNR robust regime, the angle Δ*θ*_1_ between the dominant eigenvector **v**_1_ and **D**_*τ*_ **s** is relatively small, so that cos (Δ*θ*_1_) ≈ 1 and cos (Δ*θ*_*i>*1_) ≈ 0 (due to the orthogonality of the eigenvectors). In addition, for moderate values of CFA the spread of the eigenvalues will also be moderate, and the direction of the optimal weight **w**_*dopt*_ will be primarily determined by the first eigenvector, so that the angle between **w**_*dopt*_ and **w**_*BS*_ is largely determined by Δ*θ*_1_. However, as CFA approaches 1.0, the non-dominant eigenvalues become smaller and the inverses of the smaller eigenvalues 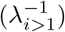 will increase such that the corresponding eigenvectors (which are orthogonal to the dominant eigenvector) will play a greater role in the composition of the optimal weight vector, moving it away from the dominant eigenvector. This will lead to an increase in the angle between **w**_*dopt*_ and the weight variants. A similar argument holds for the tSNR optimal weight vector and variants. An example of the effect is shown in Figure 8 where there is a greater spread of the weight vectors for the voxel with a higher CFA of 0.98 and a ratio of 109.7 between the maximum and minimum eigenvalues of (panel h) as compared to the voxel with a lower CFA of 0.83 and max-to-min eigenvalue ratio of 7.6 (panel g).

## 5. Discussion

We have presented a geometric view to characterize the weight dependence of tSNR and dCNR in multi-echo fMRI. Using this view, we have described three major regimes of behavior: a tSNR robust regime, a dCNR robust regime, and a within-type robust regime. The tSNR and dCNR robust regimes are characterized by covariance matrices with a dominant eigenvector. In the tSNR robust regime there is a high degree of similarity between the dominant eigenvector and the vector 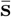 of signal means, whereas in the dCNR robust regime there is a high degree of similarity between the dominant eigenvector and the vector 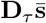 of TE-weighted signal means. Since 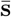 and 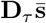 are generally not well-aligned, there is a fundamental trade-off observed within both the tSNR and dCNR robust regimes. When one of the metrics (e.g. dCNR) is relatively robust to the choice of the weight vector, the other metric (e.g. tSNR) will tend to exhibit a strong dependence on the weights. In the within-type regime, the covariance matrix does not have a dominant eigenvector and is similar in structure to the identity matrix. Within this regime, both metrics exhibit nearly optimal performance when using variants of their respective optimal weight vector. In addition, the reduced performance when using other variants is well approximated by the cosine of the angle between 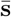 and 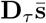. Finally, we note that for the within-type regime both the matched filter solution proposed in Posse et al. (1999) and the BOLD Sensitivity weight solution examined in Kettinger et al. (2016) approach the dCNR optimal solution, and the exponential weighting considered by Posse et al. (1999) approaches the tSNR optimal solution.

The basic behavior in each regime was first demonstrated in detail using a representative voxel for which the covariance matrix exhibited high cosine similarity to the relevant matrix (**A** for tSNR robust, **B** for dCNR robust, the identity matrix **I** for within-type robust). We then examined the brain-wide performance by assigning each voxel in the brain to the regime for which its covariance matrix showed the highest similarity to the relevant matrix (i.e. **A, B** or **I**). As this method of assigning voxels did not place constraints on the minimum allowable cosine similarity value, each regime contained voxels spanning a range of cosine similarity values. For each regime, the brain-wide normalized tSNR and dCNR distributions were characterized by strong central modes with median values that demonstrated the same tradeoffs observed for the corresponding representative voxel. In addition, the distributions exhibited long tails that were characterized within the framework of the geometric view. First, within each regime, voxels with smaller cosine similarity values deviated more from the conditions required for robust behavior. Furthermore, for the tSNR and dCNR robust regimes, low metric values were observed for voxels with high CFA values even when the cosine similarity values were relatively high. This effect reflected the occurrence of more pronounced minimums in the denominator surface and an increase in the angular spread between the optimal weight vector and the weight variants.

In the tSNR robust regime, the similarity between 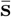 and the dominant eigenvector of the covariance matrix indicates that the noise in each echo scales with the mean signal and may be considered roughly “signal-like” (Gowland and Bowtell, 2007). In addition, since the covariance matrix can be well-approximated as a rank one matrix, the noise correlation between different echoes will be proportional to the product of the signal means of the echoes. It is important to note that the term “signal-like” does not refer to task-activated BOLD signals, but rather to signal fluctuations (e.g. due to physiological activity and scanner imperfections) that scale with the mean MRI signal level and do not demonstrate an echo time dependence (Krüger and Glover, 2001). For the dCNR robust regime, the similarity between 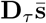 and the dominant eigenvector of the covariance matrix indicates that the noise in each echo scales with the expected BOLD signal change and may be considered roughly “BOLD-like” (Gowland and Bowtell, 2007). Furthermore, the noise correlation between different echoes will be proportional to the product of the expected BOLD signal changes of the echoes. Finally, in the within-type robust regime, the similarity between the covariance matrix and the identity matrix indicates that the noise is largely uncorrelated between echoes with a variance that is roughly independent of the echo time. These characteristics indicate that thermal noise is dominant in this regime (Posse et al., 1999; Poser et al., 2006; Gowland and Bowtell, 2007). Taken together, these observations suggest that tSNR will tend to be fairly robust to the choice of weights in brain regions where the noise is either “signal-like” or thermal-noise dominated, but less robust in regions where the noise is “BOLD-like.” On the other hand, dCNR will be fairly robust in regions where the noise is either “BOLD-like” or thermal-noise dominated, and less robust in regions where the noise is “signal-like.”

At noted in the Theory section, for task-related fMRI, tSNR and dCNR are proportional to CNR for the single-echo and multi-echo cases, respectively. In addition, for a single echo, dCNR = TE · tSNR. However, in general, the relation between tSNR and dCNR for multi-echo fMRI is not straightforward and thus the relation between tSNR and CNR is also not straightforward when there is more than one echo. For resting-state fMRI, tSNR and dCNR exhibit the same functional dependence on rSNR for the single-echo and multi-echo cases, respectively, with a linear dependence when rSNR ≪ 1. Here again, the complex relation between tSNR and dCNR means that relation between tSNR and rSNR is not straightforward when there is more than one echo. Given these observations, researchers who are currently using tSNR may want to consider also computing dCNR as a metric that is more directly related to CNR and rSNR for multi-echo acquisitions.

For the tSNR robust and within-type robust regimes, the representative voxels consisted of cerebrospinal fluid (CSF) and white matter (WM), respectively, both with partial volume fraction of 1.0. On the other hand, for the dCNR robust regime, the representative voxel consisted of slightly more gray matter (GM, fraction = 0.54) than CSF (fraction = 0.46). For the brain-wide data, the highest mean volume fractions in the tSNR robust, dCNR robust, and within-type regimes were observed for CSF, GM, and WM, respectively. The relation between operating regime and tissue composition will in general depend on the regional variations in the relative strength of signal fluctuations in the different tissue types. For example, a voxel comprised of roughly equal fractions of GM and CSF will tend to belong to the dCNR robust regime if it lies in a region (e.g. far from large vessels) in which the intrinsic BOLD-like fluctuations in GM are much greater than the signal-like fluctuations in CSF, but will tend to belong to the tSNR robust regime if it is in a region (e.g. near a large pulsatile blood vessel) where the signal-like CSF fluctuations outstrip the intrinsic BOLD fluctuations. In addition, the relation between operating regime and tissue composition will likely vary with the details of the acquisition used, such as spatial resolution, the choice of echo times, and magnetic field strength, as well as additional factors such as magnetic field inhomogeneities. For example, a decrease in voxel volume is expected to lead to a reduction in the ratio of physiological noise to thermal noise fluctuations (Triantafyllou et al., 2005; Bodurka et al., 2007) and could potentially nudge a gray matter voxel away from the dCNR robust regime towards the within-type regime. In acquisitions with pronounced magnetic field inhomogeneities, signal decay due to intra-voxel dephasing may dominate over BOLD-like changes, potentially moving a gray matter voxel from the dCNR robust regime to the tSNR robust regime. Further work to characterize the operating regimes across a range of acquisition parameters and conditions would be of interest.

The insights provided by the geometric view can serve to better guide the selection of weights, taking into account the metrics and voxels that are of primary interest. For example, if a researcher is primarily interested in dCNR and determines that the voxels that are of interest (e.g. gray matter) largely belong to the dCNR robust regime, then most weight variants will be sufficient to achieve high dCNR. However, if tSNR is the primary metric of interest while still operating in a dCNR robust regime, then a researcher may want to empirically assess the variability in tSNR across weight variants prior to selecting the weight vector to employ. On the other hand, if the voxels of interest largely belong to the within-type robust regime, a researcher may want to choose the flat weight vector since it lies between the clusters of dCNR and tSNR weight variants and thus represents a reasonable compromise that can achieve relatively high performance for both metrics.

In this work we characterized the robustness of tSNR and dCNR to the choice of weights. It is possible that tSNR and dCNR may also reflect differences in robustness to other processing choices, such as the choice of approach for physiological noise reduction. Future work is needed to explore these potential differences. For the geometric view presented in this paper, we considered a multi-echo acquisition with 3 echoes, facilitating a 3D graphical view of the weight dependence. As the arguments put forth in the Theory section do not depend on the number of echoes *N*_*E*_, the geometric view is applicable to other values of *N*_*E*_ but is difficult to visualize in higher dimensions.

As mentioned in the Introduction, both tSNR and dCNR have been used in prior studies to characterize the sensitivity of multi-echo acquisition and analysis approaches. Drawing general conclusions from the prior work can be complicated by the fact that a given study typically only uses one of these metrics. Furthermore, it is not unusual for a study to use one metric (e.g. tSNR) with a weighting scheme optimized for another metric (e.g. optimal matched-filter weighting designed to optimize CNR) (Kundu et al., 2013; Cohen et al., 2017; Heunis et al., 2021). In light of the present findings, researchers may want to consider using the framework presented here to characterize the operating regimes observed in their data and to report both tSNR and dCNR metrics. Adopting such practices could greatly facilitate the comparison of results across studies.

## 6. Data Availability Statement

The data and analysis code for this paper will be made available at https://osf.io/7awdm/ upon publication.

## 7. Acknowledgements

We thank Conan Chen, Baolian Yang, and Robert Bussell for their assistance with this work. This work was supported in part by a research grant from GE Healthcare.

## Appendix A

In this appendix we provide expanded versions of Equations 5 through 8 for the case of *N*_*E*_ = 3. The expanded version of Equation 5 is

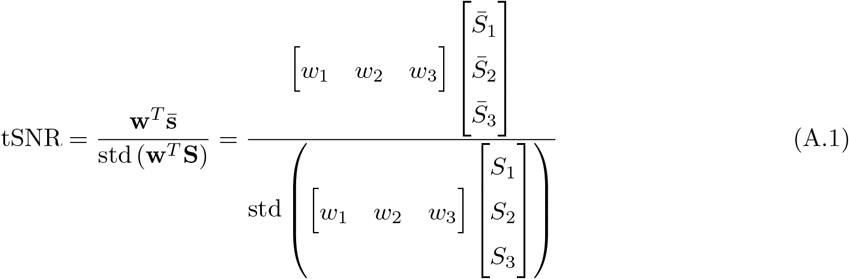

where *w*_*i*_, *S*_*i*_ and 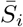 denote the weight, time series, and temporal mean, respectively, for the *i*th echo.

The expanded version of Equation 6 is

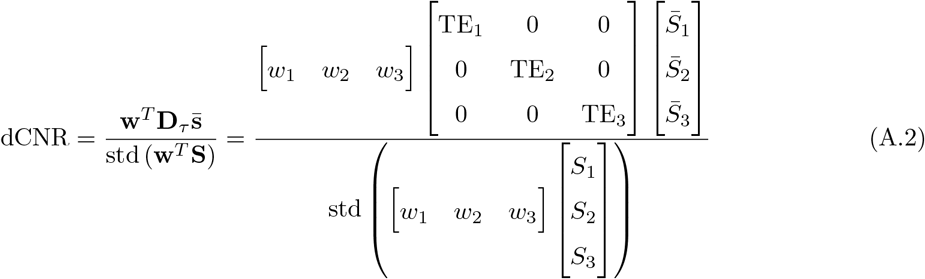

where TE_*i*_ denotes the echo time for the *i*th echo.

The expanded version of Equation 7 is

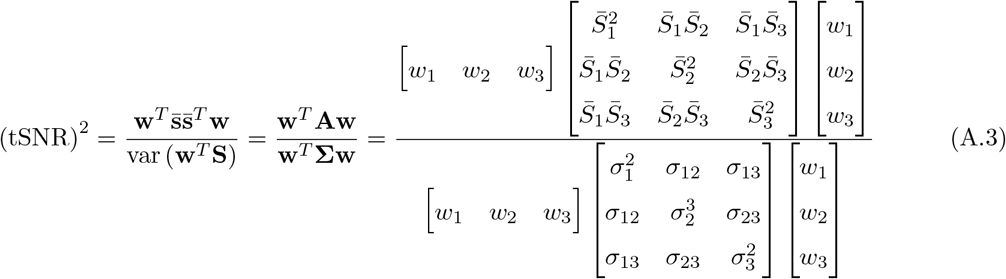

where 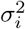 denotes the variance of the *i*th echo and *σ*_*ij*_ denotes the covariance between the *i*th and *j*th echoes with *σ*_*ij*_ = *σ*_*ji*_.

The expanded version of Equation 8 is

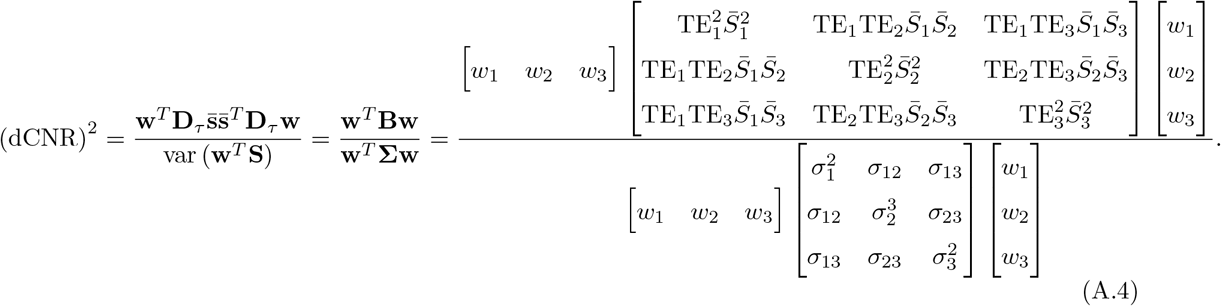

